# Microplastics dysregulate innate immunity in the SARS-CoV-2 infected lung

**DOI:** 10.1101/2023.11.19.567745

**Authors:** Cameron R. Bishop, Kexin Yan, Wilson Nguyen, Daniel J. Rawle, Bing Tang, Thibaut Larcher, Andreas Suhrbier

## Abstract

Global microplastic (MP) pollution is now well recognized, with humans and animals consuming and inhaling MPs on a daily basis. Herein we described the effects of azide-free, 1 µm polystyrene MP beads co-delivered into lungs with a SARS-CoV-2 omicron BA.5 inoculum using a mouse model of mild COVID-19. Lung virus titres and viral RNA levels were not significantly affected by MPs, with overt clinical or histopathological changes also not observed. However, RNA-Seq of infected lungs revealed that MP exposure suppressed innate immune responses at 2 days post infection (dpi) and increased pro-inflammatory signatures at 6 dpi. The cytokine profile at 6 dpi showed a significant correlation with the ‘cytokine release syndrome’ signature seen in some severe COVID-19 patients. This study adds to a growing body of literature suggesting that MPs can dysregulate inflammation in specific disease settings.

**Graphical Abstract:** *HIGHLIGHTS:*

- A single inoculation of microplastics dysregulated SARS-CoV-2 lung inflammation
- At the peak of SARS-CoV-2 infection microplastics decreased early innate responses
- Later post infection microplastics promoted a “cytokine release syndrome” signature
- A key mechanism may involve the inhibition of the phagocytosis of infected cells
- Azide-free microplastics were used, with no elevated ROS responses identified Postulated mechanisms whereby microplastics might decrease the proinflammatory responses 2 days after SARS-CoV-2 infection, yet promote the proinflammatory ‘cytokine release syndrome’ signature at 6 days post infection.

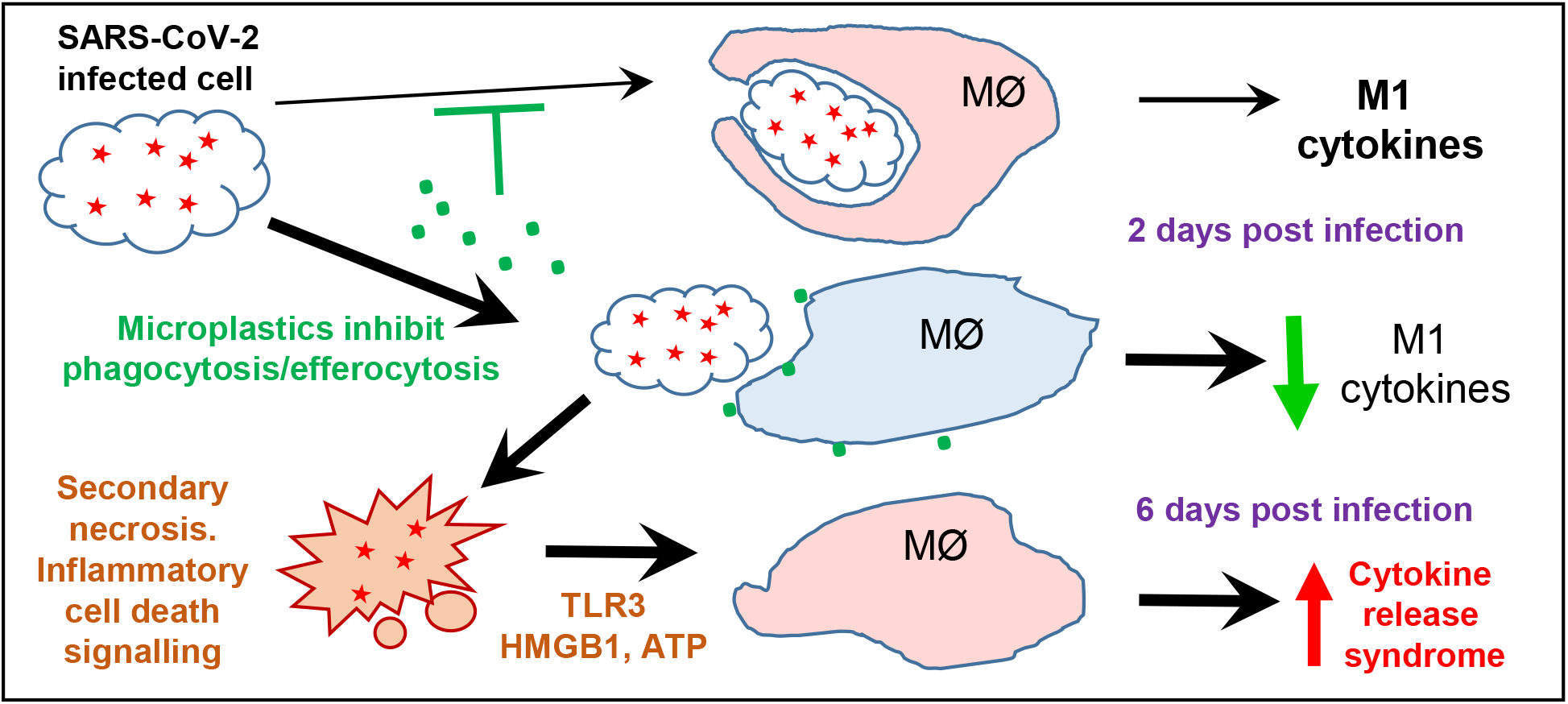

## 1. Introduction

Global plastic production has grown exponentially, nearly 475 million tons were produced in 2021, and this is set to climb to ≈ 550 million tons in 2026. The latter part of the Anthropocene can now be referred to as the Plasticene (Rangel-Buitrago et al., 2022), with the plastic debris providing a new ecological niche known as the plastisphere (Du Toit, 2022). Plastics degrade into microplastics (MPs), with MP contamination of our environment now well described, arising from both use of items manufactured from plastic, as well as poor disposal of plastic waste resulting in plastic pollution (Kutralam-Muniasamy et al., 2023; Le et al., 2023; Priya et al., 2023). Less clear is what the implications are for the health of human populations, with considerable speculation available (Blackburn et al., 2022; Jones et al., 2023; Koelmans et al., 2022; Vethaak et al., 2021), but a noteworthy paucity of compelling direct evidence for detrimental human clinical outcomes caused by MP exposure. Nevertheless, some clinical data is emerging. For instance, a positive correlation was reported between faecal MP concentration and the severity of inflammatory bowel disease (Yan et al., 2022b). More MPs were present in the lungs of paediatric patients with severe community acquired pneumonia (CAP) when compared with those with non-severe CAP (Chen et al., 2023). MPs were found in cirrhotic liver tissue, but not in liver samples from individuals without underlying liver disease (Horvatits et al., 2022). Unfortunately, whether the different MP levels identified in these studies were causative or a consequence of the disease, remains unclear. More robust evidence for causation is seen for occupational diseases in workers from synthetic textile, flock and (poly)vinyl chloride industries, following exposure to chronic high doses of airborne MPs (Prata, 2018; Zuri et al., 2023). For instance, chronic interstitial pneumonitis and breathing difficulties have been associated with workplace exposure to nylon flock (Eschenbacher et al., 1999; Washko et al., 2000).

Even outside heavily contaminated industrial settings, humans are exposed to airborne MPs at home, at the office and outdoors (Liu et al., 2019a; Perera et al., 2023; Romarate et al., 2023), with widespread reports of MPs identified in human lungs (Baeza-Martinez et al., 2022; Chen et al., 2023; Jenner et al., 2022; Jiang et al., 2022; Qiu et al., 2023) and sputum (Huang et al., 2022). Estimates from several studies for MP inhalation in the home were 0.3-2.5 µg/kg/d (Soltani et al., 2021). Using indoor dust measurements from 12 countries, the median MP intake was calculated to be 0.36-150 µg/kg/d, with adult intake 10 fold lower than infants (Zhang et al., 2020). An estimated MP inhalation of 6.5-8.97 µg/kg/day is reported to be derived from human exposure models (Danso et al., 2022; Kannan et al., 2021; Wang et al., 2021), although where this data came from or how it was derived is unclear.

Rodent studies seeking to assess the effects of MP inhalation, introduce MPs into the lungs via intranasal or intratracheal inoculation. Unfortunately, as for many such studies on MP exposure, the doses used were often unrealistically high (Zolotova et al., 2022). Doses were, for instance, 40 mg/kg/d for 21 days of carboxy-modified 1-5 and 10-20 µm polystyrene beads (Cao et al., 2023), 5 mg/kg/d for 2 weeks of three types of MPs (Danso et al., 2022), 5 mg/kg 3 times per week of 100 nm amino-modified polystyrene beads (Wu et al., 2023), 4 mg/kg (assuming 25 g mouse) 5 µm and 99 nm polystyrene beads every other day for five weeks (Zha et al., 2023), or 1.25 and 6.25 mg/kg 3 times per week of 5 µm polystyrene beads for three weeks (Li et al., 2022). Such doses are 1 to 2 orders of magnitude higher than even the highest estimates for humans. Another serious confounding issue is the potential presence of highly toxic preservatives such as azide, which are commonly added to commercial polystyrene bead products (Pikuda et al., 2019). Such chemicals would provide both acute toxicity if the beads are not washed, and would also slowly leach out of the MPs *in vivo,* imbuing the MPs with artificial toxicity. Azide inhibits mitochondrial cytochrome C oxidase (the terminal complex of eukaryotic oxidative phosphorylation) and catalase. Catalase protects cells against oxidative damage by reactive oxygen species (ROS). ROS associated toxicity is frequently reported for MP exposure (Hu et al., 2020; Subaramaniyam et al., 2023) and may thus simply represent an azide artefact.

An important recent observation is that MPs can be recognized by Tim4, a receptor that plays an essential role in binding and phagocytosing apoptotic cells (a process known as efferocytosis), with MPs able to inhibit efferocytosis by macrophages *in vitro* (Kuroiwa et al., 2023). Azide free, 0.8 µm beads (Sigma-Aldrich, LB8) were used for many of the experiments, with no toxicity or induction of inflammation observed for macrophages treated with MPs *in vitro* (Kuroiwa et al., 2023). Efferocytosis promotes anti-inflammatory activities and inflammation resolution, with efferocytosis failure or defects resulting in apoptotic cells undergoing secondary necrosis, which is generally pro-inflammatory *in vivo* (Mehrotra et al., 2022; Schilperoort et al., 2023).

We have previously shown that consumption of azide-free 1 µm polystyrene beads had minimal overt effects by themselves, but after infection with the arthritogenic chikungunya virus, intestinal MPs caused a significant prolongation of the ensuing viral inflammatory arthritis (Rawle et al., 2022). Given the recent SARS-CoV-2 pandemic and the aforementioned estimates on human MP inhalation, we sought herein to examine whether the presence of MPs in the lung would affect the outcome of SARS-CoV-2 infection and COVID-19 disease in a recently established non-lethal mild transgenic mouse model, where the virus receptor, human angiotensin converting enzyme 2 (ACE2) is expressed from the mouse ACE2 promoter (Bishop et al., 2022). The inflammatory response and the cytokine responses associated with COVID-19 are well described (Al-Nesf et al., 2022; Maity et al., 2023), and are largely recapitulated in mouse models (Bishop et al., 2022; Liu et al., 2023; Rawle et al., 2021). We also used a recent SARS-CoV-2 variant of concern, omicron BA.5 (Yan et al., 2022a), with omicron variants currently the dominate SARS-CoV-2 viruses infecting human populations (Our_World_in_Data, 2023). Using this mouse model system and RNA-Seq, we illustrate that a single co-inoculation of MPs and virus into the lungs dysregulates the innate inflammatory response to SARS-CoV-2, moving the cytokine response profile towards a “cytokine release syndrome” signature.

## 2. Materials and Methods

### 2.1 Ethics statements and PC3/BSL3 certifications

All mouse work was conducted in accordance with the “Australian code for the care and use of animals for scientific purposes” as defined by the National Health and Medical Research Council of Australia. Mouse work was approved by the QIMR Berghofer Medical Research Institute animal ethics committee (P3600), with infectious SARS-CoV-2 work conducted in a PC3 (Bio-Safety Level 3) facility at the QIMR Berghofer MRI (Australian Department of Agriculture, Water and the Environment certification Q2326 and Office of the Gene Technology Regulator certification 3445). Breeding and use of GM mice was approved under a Notifiable Low Risk Dealing (NLRD) Identifier: NLRD_Suhrbier_Oct2020: NLRD 1.1(a). Mice were euthanized using carbon dioxide.

Collection of nasal swabs from consented COVID-19 patients (to isolate circulating SARS-CoV-2 variants) was approved by the QIMR Berghofer Medical Research Institute Human Research Ethics Committee (P3600).

### 2.2 The SARS-CoV-2 omicron BA.5 virus isolate

The omicron BA.5 isolate, SARS-CoV-2_QIMR03_ (SARS-CoV-2/human/AUS/QIMR03/2022) belongs to the BE.1 sublineage (GenBank: OP604184.1) and was obtained at QIMR Berghofer MRI from nasal swabs from a consented COVID-19 patient (Stewart et al., 2023). Virus stocks were propagated in Vero E6 cells, stocks and tissue culture supernatants were checked for endotoxin (Hirata et al., 2018; Johnson et al., 2005) and mycoplasma (MycoAlert, Lonza) (La Linn et al., 1995). Virus titres were determined by CCID_50_ assays (Yan et al., 2021).

### 2.3 mACE2-hACE2 mice and infection

mACE2-hACE2 mice (Bao et al., 2020) were generated as described (Bishop et al., 2022) by Monash Genome Modification Platform (MGMP), Monash University and are freely available through Phenomics Australia (MGMP code ET26). The strain was initially maintained in-house as heterozygotes by backcrossing to C57BL/6J mice. Heterozygotes were inter-crossed to generate a homozygous mACE2-hACE2 transgenic mouse line. Genotyping was undertaken by digital droplet PCR (MGMP) to distinguish homozygotes from heterozygotes; hACE2 primers 5ʹ-CCAGATGTACCCTCTGCAAG-3ʹ/5ʹ-TCGTGTTCAGGATGGTGTTC-3ʹ, probe 6-carboxyfluorescein-5ʹ-GCTCCAGCTGCAGGCTCTCCAGCA-3ʹ-ZEN/IowaBlack; RPP30 reference primers CTTTGAACTTGTCTATGGTCCT/GCATCAAATTGAGGGCATTG, probe hexachlorofluorescein-TGTGTACCTTCTCATCGTTGCATC-ZEN/IowaBlack. A Bio-Rad QX200 ddPCR droplet generator was used to generate droplets, amplified products were analysed by QX200 droplet reader, and copy numbers were determined using QuantaSoft Analysis Pro Version 1.0 (Bio-Rad, USA). After 2 inter-crossing of homozygotes, all offspring were homozygotes, and a homozygous line was established.

Mice were infected as described (Dumenil et al., 2023), briefly, female mice (≈10-20 weeks of age) received intrapulmonary infections delivered via the intranasal route with 5×10^4^ CCID_50_ of virus in 50 μl RPMI 1640, while under light anaesthesia. Each group of mice within an experiment had a similar age range and distribution, with the mean age for each group not differing by more than 1 week. Mice were weighed and overt disease symptoms scored as described (Stewart et al., 2023). Mice were euthanized using CO_2_, and tissue titres determined using CCID_50_ assays and Vero E6 cells (Yan et al., 2021).

### 2.4 Microplastics (MPs)

The MPs comprised internally dye loaded, Fluoresbrite® yellow-green polystyrene-based microspheres with a diameter of 1 µm (Cat# 17154-10) purchased from Polysciences as 2.5% w/v in a sterile aqueous suspension without sodium azide. The zeta potential of ≈1 µm polystyrene beads has been estimated to be ≈ -20 mV (Delgado et al., 1986). The rationale for choosing these MPs has been described previously (Rawle et al., 2022); briefly, polystyrene MPs are frequently found in the environment, surface labelled microspheres have altered surface characteristics not recapitulate by MPs in the environment, and the 1 μm size is approximately the size of a bacteria, with bacteria routinely phagocytosed by macrophages. MPs were diluted in PBS and administered alone or together with the viral inoculum in a single dose of 1 µg of MPs per mouse (≈40 µg/kg); total inoculated volume was always 50 µl per mouse.

### 2.5 MP visualisation and quantitation in lung tissues

Lungs were fixed in 10% formalin for 2-3 days, tapped dry and embedded in O.C.T. (Tissue-Tek, Qiagen), with ≈ 7 µm cryosections, under a glass coverslip, viewed by fluorescent microscopy. MPs were counted by eye.

Lungs were weighted, manually chopped using scissors and digested in ammonium sulphate (50 mM), SDS (5 mg/ml) and proteinase K (1 mg/ml) overnight at 37°C as described (Walczak et al., 2015). Digested suspensions were viewed by fluorescent microscope and a haemocytometer, with fluorescent MPs counted by eye.

### 2.6 Histology and immunohistochemistry

Histology and immunohistochemistry were undertaken as described (Yan et al., 2022a), with lungs fixed in formalin embedded in paraffin, sections stained by H&E and slides scanned by Aperio AT Turbo (Aperio, Vista, CA, USA). Image analysis (nuclear/cytoplasmic staining ratios) was undertaken using Positive Pixel Count v9 algorithm. White space analysis was undertaken using QuPath v0.2.3.

Immunohistochemistry (IHC) was undertaken as described (Morgan et al., 2022) using the macrophage/monocyte monoclonal antibody F4/80 (Abcam, Cambridge, MA) and colour developed using NovaRed (Vector Laboratories, Newark, CA, USA).

### 2.7 RNA-Seq and bioinformatic analyses

RNA-Seq and bioinformatic analyses were undertaken as described (Bishop et al., 2022; Dumenil et al., 2023). Briefly, mouse lung tissues were harvested into RNAlater, RNA was extracted using TRIzol (Life Technologies), and RNA concentration and quality measured using TapeStation D1kTapeScreen assay (Agilent). cDNA libraries were generated using Illumina TruSeq Stranded mRNA library prep kit and sequencing performed in-house using Illumina Nextseq 550 platform (75-base paired end reads). Processed reads were aligned to GRCm39 vM26 (mouse genome) and the BA.5 genome using STAR aligner. Gene expression was calculated using RSEM and EdgeR. For the BA.5+MP vs. BA.5-MP data sets a term for viral reads was introduced into Edge R to minimize the effects of viral loads on significance and fold change in the mRNA expression data. A filter was applied to the count matrix of counts per million (cpm) >1 for any given gene in at least 5 samples.

Differentially expressed genes (DEGs) were analysed using Ingenuity Pathway Analysis (IPA) (QIAGEN). Whole gene lists ranked by fold change were interrogated using Gene Set Enrichment Analyses (GSEA v4.0.3) (Broad Institute, UCSanDiego) using the “GSEAPreranked” module. Gene sets were obtained from the complete Molecular Signatures Database (MSigDB) v7.2 (31,120 gene sets) (msigdb.v7.2.symbols.gmt), Blood Transcription modules (BTMs), and Xue at al., 2014 (Xue et al., 2014). Differences in specific cell types were analysed using cell-type deconvolution, SpatialDecon (Danaher et al., 2022), with cell-type expression matrices obtained from Yoshida et al. 2019 (Yoshida et al., 2019) or the NanoString Cell Profile Library, either Mouse/Adult/Lung_MCA (Mouse cell atlas) or Mouse/Adult/ImmuneAtlas_ImmGen_cellfamily (Immune cell family).

Kraken metagenomic sequence classification was undertaken as described (Hazlewood et al., 2021).

### 2.8 Statistics

Statistical analyses of experimental data were performed using IBM SPSS Statistics for Windows, Version 19.0 (IBM Corp., Armonk, NY, USA). The t-test was used when the difference in variances was <4 (determined using Data Analysis ToolPak in Excel), skewness was >-2 and kurtosis was <2 (determined using SPSS). Otherwise, the non-parametric Kolmogorov-Smirnov test was used.

## 3. Results

### 3.1 MP inoculation into mouse lungs promotes mild inflammation on day 2

Delivery of MPs into mouse lungs generally involves delivery of MPs in solution into the lungs via the intranasal (i.n.) route (Cao et al., 2023; Danso et al., 2022; Li et al., 2022; Wu et al., 2023; Zha et al., 2023), as mimicking MP dust inhalation is currently technically, logistically and ethical difficult in a laboratory setting. Some of the challenges include consistent airborne MP dosing, staff safety considerations, and mouse eye irritation, respectively.

To assess the effects of MPs on uninfected lungs, a single dose of 1 µg of MPs per mouse (≈40 µg/kg) of azide-free, 1 µm diameter, fluorescent dye loaded, polystyrene beads (density 0.26 g/ml) was delivered in 50 µl of PBS via the i.n. route into the lungs of lightly anesthetized female C57BL/6J mice. Control mice receiving 50 µl of PBS. The anaesthesia prevents the mouse sneezing or coughing out the introduced material, but is light enough for the mouse to retain largely normal breathing, promoting deep lung delivery of the material; this is the same method used to infect mice with SARS-CoV-2 (Amarilla et al., 2021; Bishop et al., 2022; Dumenil et al., 2023; Mills et al., 2021; Yan et al., 2022a). The mice were observed daily and no overt clinical signs were observed. Mice were euthanized on day 2 and 6 post MP inoculation, and lungs analysed by RNA-Seq, with bioinformatic treatments comparing +MP day 2 vs. PBS day 2 and +MP day 6 vs. PBS day 6 (Fig. 1; Supplementary Table 1 and 2).

**Figure 1.**
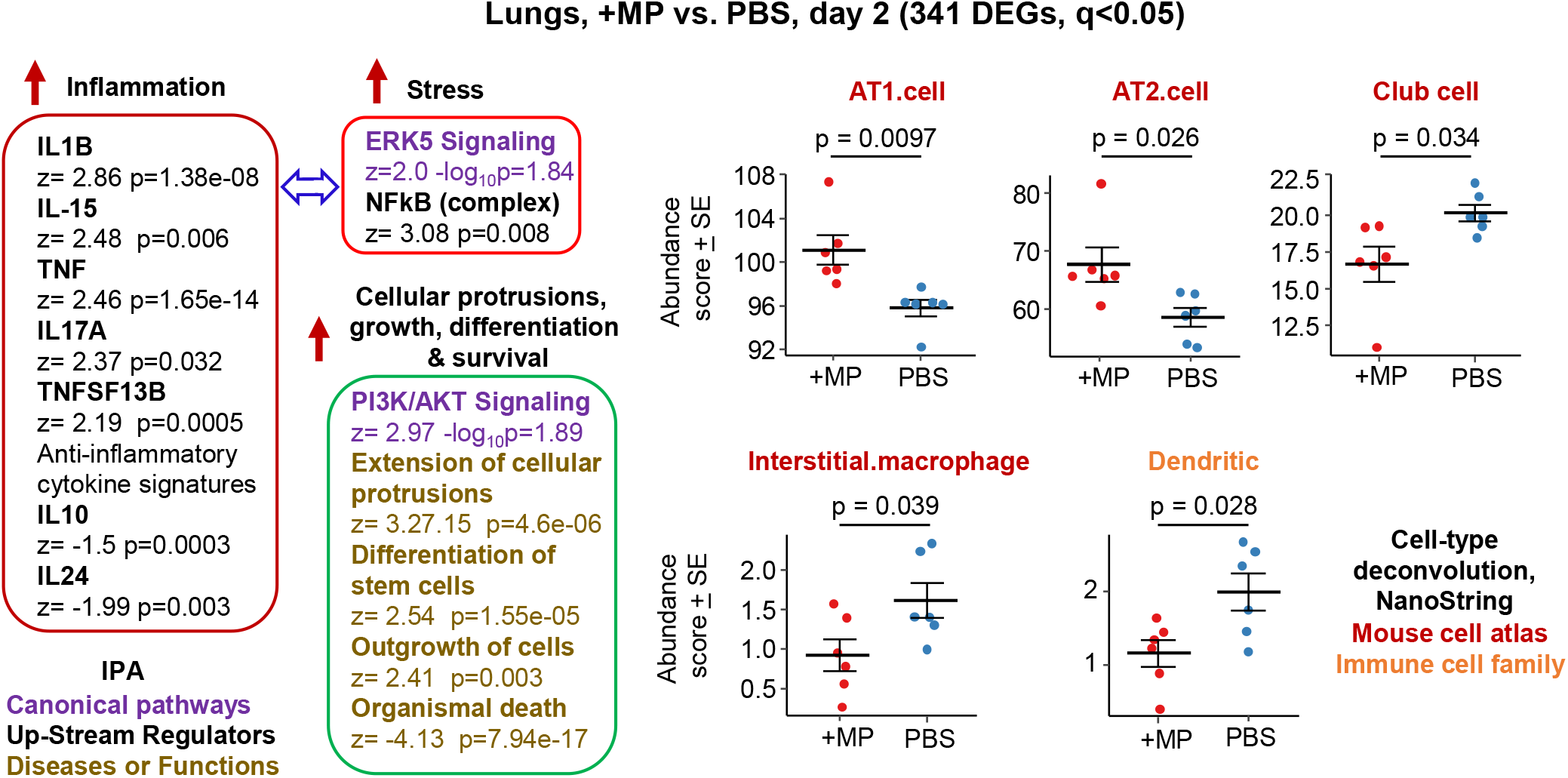
Bioinformatic analyses of RNA-Seq data for lungs inoculated with MP. C57BL/6 mice were inoculated with MP or PBS into the lungs via the intranasal route and at 2 days post inoculation lungs were harvested and analysed by RNA-Seq. 341 DEGs were identified and analysed by IPA, with selected annotations shown, full data sets are provided in Supplementary Table 1. The whole gene list (Supplementary Table 1) was also analysed by cell type deconvolution using cell-type expression matrices obtained from NanoString; statistics by t tests.

RNA-Seq for +MP vs. PBS day 2 identified 341 differentially expressed genes (DEGs) at q (FDR) <0.05 (Supplementary Table 1). Fold change (FC) was low overall, with only 9 genes showing FC>2 (log2 FC>1) and only 4 of these had a mean normalized counts per million (cpm) >10 across all samples (Supplementary Table 1). Ingenuity Pathway Analysis (IPA) indicated a mild pro-inflammatory response, with IL-1β the highest Cytokine UpStream Regulator (USR) by z-score (Fig. 1, Inflammation; Supplementary Table 1). Some classical stress-associated signatures were also evident (Fig. 1, Stress). A series of annotations associated with cellular protrusions, proliferation, differentiation and survival were identified as the top annotations (by z score) by IPA Diseases and Functions (Fig. 1), with the largest negative z scores associated with growth failure or death (Supplementary Table 1).

Differences in specific cell types were analysed using cell-type deconvolution, SpatialDecon (Danaher et al., 2022), with cell-type expression matrices obtained from Yoshida et al. 2019 (Yoshida et al., 2019) or the NanoString Cell Profile Library, either Mouse/Adult/Lung_MCA (Mouse cell atlas) or Mouse/Adult/ImmuneAtlas_ImmGen_cellfamily (Immune cell family) (Supplementary Table 1). These analyses suggested that the cells that were expanding and/or more transcriptionally active in the +MP group were AT1 and AT2 type I and II alveolar epithelial cells (Fig. 1, AT1, AT2). The positive “Normalised Enrichment Score” (NES) for “Differentiation of stem cells”, may be associated with reduced abundance of Club cells (formally known as Clara cells), which are regional progenitor cells that repair bronchiolar epithelium in response to lung damage. Also identified was a reduced abundance of interstitial macrophages, cells that are associated with nerves and airways and appear to have important immunoregulatory roles (Aegerter et al., 2022; Ural et al., 2020). A reduction in interstitial macrophages may also be consistent with the reductions in anti-inflammatory cytokines (Fig. 1, IL-10 and IL-24) (Ural et al., 2020). Dendritic cell abundance was also down in the +MP group, perhaps consistent with mobilization to lymph/lymph nodes in response to inflammation (Prow et al., 2010).

In summary, MP inoculation into lungs generated a transcriptional signature indicating a mild inflammatory response, and mild increases in abundance scores (likely proliferation) for lung epithelial cells, and reductions in abundance scores for interstitial macrophages and dendritic cells. Signatures associated with MP induction of ROS were not identified; IPA Disease and Functions annotation actually providing a slightly negative z-score for “Production of reactive oxygen species” (Supplementary Table 1).

### 3.2. Effects of a single MP inoculation into lungs are largely resolved within 6 days

Harvesting of lungs on day 6 after MP or PBS inoculation and comparing the transcriptome by RNA-Seq (+MP day 6 vs. PBS day 6) identified only 7 DEGs, all with low fold change (Supplementary Table 2). This argues that the lung response to a single exposure to MPs is largely resolved within 6 days. GSEAs using MSigDB gene sets provided a series of significant annotations with high positive NES associated with epithelial cells (e.g. Epithelial differentiation), suggesting some lung repair activities were still underway on day 6 (Supplementary Table 2).

### 3.3 The mACE2-hACE2 omicron BA.5 mouse model of SARS-CoV-2 infection and disease

The best described mouse model for SARS-CoV-2 infection is the K18-hACE2 model which expresses the SARS-CoV-2 receptor, human ACE2 (hACE2), from the keratin 18 promoter (K18). This model is generally lethal within several days, as it usually leads to brain infection and ensuing weight loss that reaches ethically defined criteria for euthanasia (Dumenil et al., 2023; Mills et al., 2021; Stewart et al., 2023). A non-lethal, less severe model is the mACE2-hACE2 model, wherein hACE2 is expressed from the mouse ACE2 promoter (mACE2) (Bao et al., 2020). We generated such a transgenic mouse on a pure C57BL/6J background by microinjection of the mACE2-hACE2 transgene into the pronucleus of C57BL/6J zygotes at the pronuclei stage (Bishop et al., 2022) and generated a homozygous mACE2-hACE2 transgenic mouse line (see Materials and Methods). Given MP-mediated effects were generally mild (Fig. 1; Supplementary Table 1), we chose this mouse model so that any MP-mediated perturbations might be more readily detected.

Although infection of mACE2-hACE2 mice with an original strain isolate has been described (Bao et al., 2020; Bishop et al., 2022), infection of homozygous mACE2-hACE2 mice with an omicron BA.5 isolate has not. The latter did not result in significant weight loss (data not shown), with lung histology at 6 days post infection (dpi) showing a series of histopathological features that have been described previously in COVID-19 mouse models (Amarilla et al., 2021; Bao et al., 2020; Rawle et al., 2021) (Supplementary Fig. 1). Lesions were less severe when compared with those seen after infection of K18-hACE2 mice with an original strain isolate (Arce et al., 2021; Dumenil et al., 2023; Winkler et al., 2020), although significant loss of white space (unstained air-spaces) in H&E stained lung sections (indicating lung consolidation) (Amarilla et al., 2021; Yan et al., 2022a), was also seen in this model (Supplementary Fig. 2).

Infection of mACE2-hACE2 mice with BA.5 (BA.5 vs. PBS) was analysed by RNA-Seq, with lungs harvested on 2 dpi (peak viral load) and 6 dpi (peak lung pathology) and were compared with mock infected lungs (PBS) harvested on days 2 and 6, respectively (Supplementary Tables 3 & 4). When the lung cytokine response signatures (IPA Cytokine USR z-scores) from BA.5-infected mACE2-hACE2 mice, were compared with those from K18-hACE2 mice infected with an original strain isolate (Bishop et al., 2022), a highly significant correlation emerged for 2 dpi (Supplementary Fig. 3a, Pearson correlation p=1.95×10^-28^ r=0.8). The correlation for 5/6 dpi was less significant (Supplementary Fig. 3b, Pearson correlation p=0.0005 r=0.33), likely reflecting the increased disease severity in K18-hACE2 mice, with more cytokine USR annotations with higher z scores seen in the K18-hACE2 model.

IPA Diseases and Function annotations for BA.5 infected lungs for mACE2-hACE2 mice (BA.5 vs. PBS) pertinent to the analyses below include, top annotations for leukopoiesis/haematopoiesis, phagocytosis (engulfment of cells) and apoptosis & necroptosis (Supplementary Tables 3 & 4).

### 3.4. MP clearance from lungs is slower in BA.5 infected mice

To evaluate the effects of SARS-CoV-2 infection on MP clearance from lungs, mACE2-hACE2 mice were given a single inoculum of 50 µl containing both BA.5 (5×10^4^ CCID_50_) and 1 µg (≈40 µg/kg) of azide free, 1 µm diameter, florescent dye loaded, polystyrene beads (BA.5+MP). The inoculum was delivered into the lungs via the intranasal route using the same procedure used herein and generally to infect mice with SARS-CoV-2 (Amarilla et al., 2021; Dumenil et al., 2023; Guimond et al., 2022). Control mice were not infected, but received the same 50 µl inoculum containing MP (+MP). Mixing the beads with the viral inoculum prior to infection prevented any complications that might arise from simple liquid occlusion of airways, if for, instance, the SARS-CoV-2 inoculum was given first and MPs were introduced later.

Lungs were harvested at ≈4 hrs, 24 hrs (day 2), 48 hrs and 144 hrs (day 6), were fixed in formalin, and the beads (MPs) observed by fluorescent microscopy of cryosections (Fig. 2a). Quantitation suggested reduced clearance of the MPs in BA.5-infected lungs (Fig. 2b), consistent with SARS-CoV-2-mediated disruption of the ciliary layer, which is responsible for mucociliary clearance (Dumenil et al., 2023; Robinot et al., 2021). To provide better quantitation, lungs were digested in proteinase K and dissolved in SDS, and beads counted under a hemocytometer. The same trend was observed, with significantly more beads present at 24 and 48 hrs in infected mice (Fig. 2c). Both assays systems indicated that MPs were largely cleared by day 6 (Fig. 2b,c).

**Figure 2.**
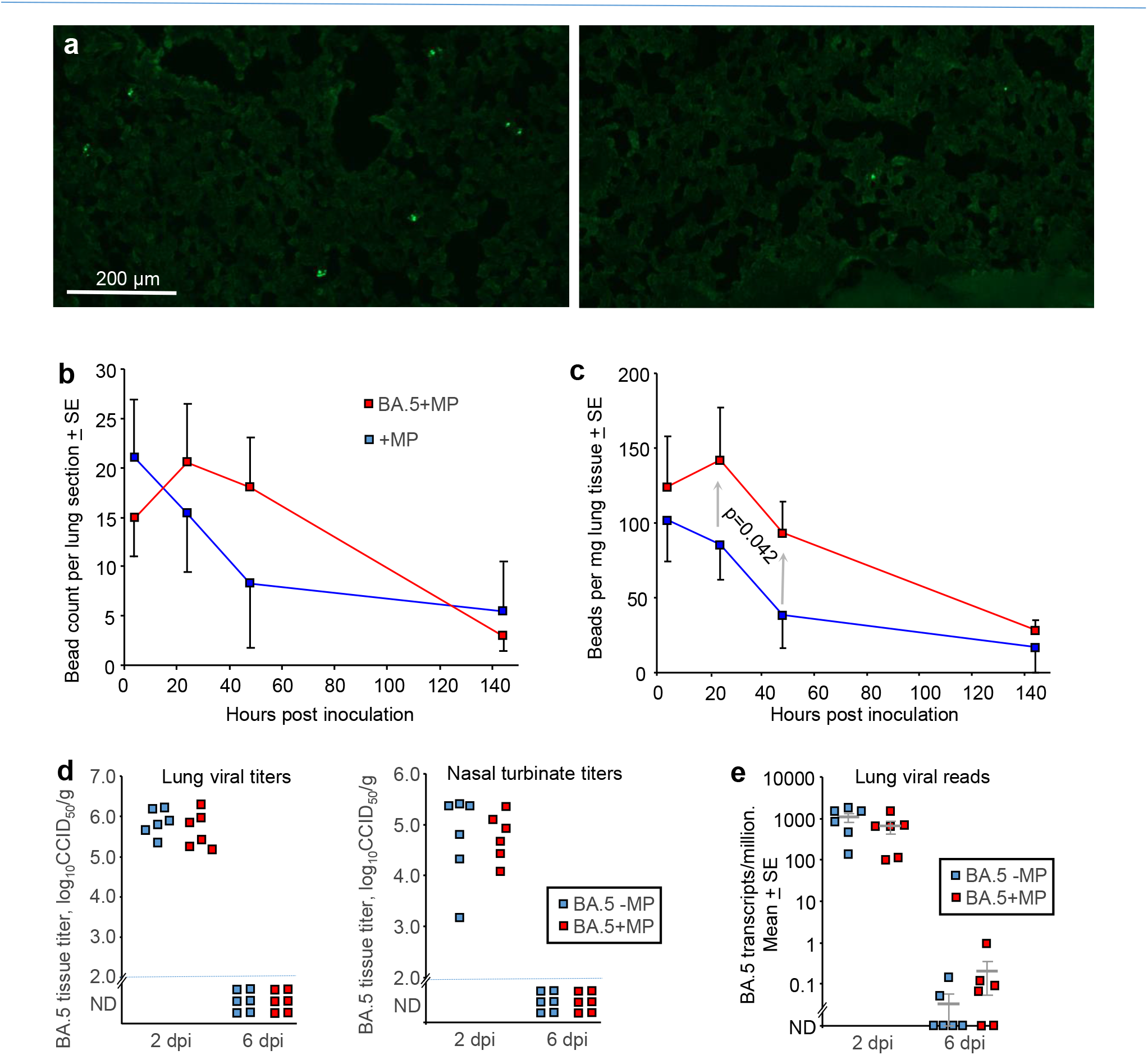
SARS-CoV-2 infection slowed MP clearance, but MP did not affect viral load. **a** mACE-hACE2 mice were inoculated with BA.5+MP or with just MP (+MP), lungs were harvested at different time points and cryosections of lungs observed by fluorescent microscopy. Examples are shown for the 4 hr (right) and 48 hr (left) time points for the +MP group. **b** Quantitation of MP counts from cryosections (n=6 mice per group for 4 hrs, n=3 for other time points). **c** Quantitation of MP counts using tissue digestion and hemocytometer. The two groups were statistically different by 2 way ANOVA, which included a term for hours post inoculation with data for 24 and 48 hrs included. **d** mACE-hACE2 mice were inoculated with BA.5+MP or with BA.5-MP (no MP) and lung and nasal turbinate tissue titers determined by CCID_50_ assays. **e** mACE-hACE2 mice were inoculated with BA.5+MP or with BA.5-MP and lung analysed by RNA-Seq. Viral read counts are shown as BA.5 transcripts per million.

### 3.5. Lung viral loads were unaffected by MPs

To evaluate the effects of MPs on SARS-COV-2 infection, mACE2-hACE2 mice were given a single inoculum of 50 µl containing both BA.5 (5×10^4^ CCID_50_) and 1 µg (≈40 µg/kg) of beads, delivered into the lungs via the intranasal route (BA.5+MP). Control mice received the same 50 µl inoculum containing BA.5, but no MP (BA.5-MP). Lungs and nasal turbinates were harvested on 2 and 6 days post infection (dpi) and tissue titres determined by CCID_50_ assays, with no significant differences in viral titres evident on 2 dpi, and below the level of detection by 6 dpi (Fig. 2d).

Lungs were also analysed by RNA-Seq and reads aligned to the viral genome, with the number of viral reads not significantly different on 2 or 6 dpi (Fig. 2e; Supplementary Tables 5 and 6). Thus overall MP inoculation had no significant effects on viral loads in the respiratory track.

### 3.6. MPs reduced innate proinflammatory signatures in BA.5 infected lungs 2 dpi

To assess the effects of MPs on the innate immune responses induced by BA.5 infection, the same groups described above (BA.5+MP vs. BA.5-MP, 2 dpi) were compared by RNA-Seq, with reads aligned to the mouse genome. As there were small non-significant differences in viral reads between the two groups (Fig. 2e, Supplementary Table 5), a term for viral reads was introduced into Edge R to minimize the effects of viral loads on significance and fold change in the mRNA expression data. Applying a q<0.05 filter, 596 DEGs were thereby identified (Fig. 3, Supplementary Table 5).

**Figure 3.**
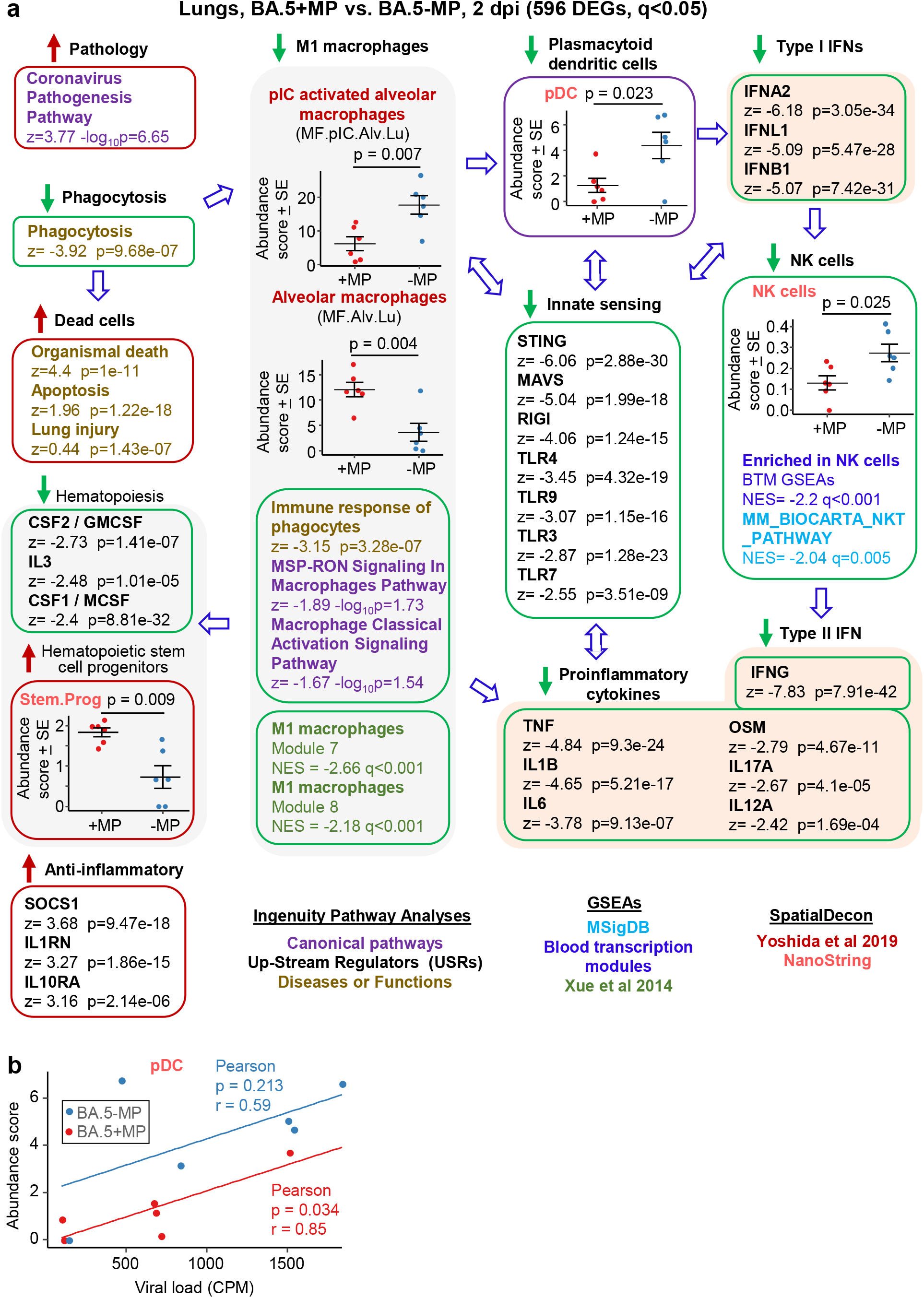
Bioinformatic analyses of RNA-Seq data for BA.5+MP vs. BA.5-MP at 2 dpi. **a** mACE-hACE2 mice were inoculated with BA.5+MP or with BA.5-MP and at 2 dpi lungs were harvested and analysed by RNA-Seq. 596 DEGs were identified and analysed by IPA, with selected annotations shown; full data sets are provided in Supplementary Table 5. The whole gene list was also analysed by GSEAs, and cell type deconvolution (SpacialDecon) using cell-type expression matrices obtained from Yoshida et al 2019 and NanoString (statistics by t tests). **b** Cell type deconvolution data for plasmacytoid dendritic cells shown in ‘a’ plotted against viral reads (counts per million). Pearson correlation significance (p) and correlation coefficient (r) provided.

The DEGs were analysed by IPA as above, and the All gene list, ranked by fold change, was used in a series of Gene Set Enrichment Analyses (GSEAs) using gene sets from the Molecular Signatures Data Base (MSigDB), Blood Transcription modules (BTM) and Xue et al., 2014 (Xue et al., 2014) (Fig. 3, Supplementary Table 5). Differences in specific cell types were analysed as above using SpatialDecon (Danaher et al., 2022), using gene expression matrices provided by NanoString Cell Profile Library, and Yoshida *et al*., 2019 (Yoshida et al., 2019) (Fig. 3, Supplementary Table 5).

The highest ranked IPA Canonical pathway was the ‘Coronavirus pathogenesis pathway’ (Fig. 3a, Pathology; Supplementary Table 5). MPs were recently reported to inhibit phagocytosis of apoptotic cells (known as efferocytosis) (Kuroiwa et al., 2023), and a top IPA Diseases and Functions annotation emerged to be ‘Phagocytosis’, with a high negative z score (Fig. 3a, Phagocytosis; Supplementary Table 5). During a viral infection, macrophages phagocytose SARS-CoV-2-infected cells and are thereby activated, and in turn mediate strong activation of plasmacytoid dendritic cells (pDC) (Garcia-Nicolas et al., 2023). The reduced phagocytosis is thus consistent with a reduction in activated M1 macrophages identified via a series of annotations (Fig. 3, M1 macrophages); specifically (i) reduced abundance of polyinosinic:polycytidylic acid (pIC) stimulated macrophages (a stimulus that mimics viral double stranded RNA); (ii) a series of IPA Canonical pathways and Diseases and Functions macrophage annotations with negative z scores; and (iii) GSEAs with negative NES and q<0.05 using M1 macrophage gene sets from Xue at al., 2014 (Xue et al., 2014) (Supplementary Table 5).

Significantly reduced numbers of pDC were identified in the BA.5+MP group by SpacialDecon using NanoString gene lists (Fig. 3, pDC, Supplementary Table 5), consistent with the reduction in M1 macrophages (Garcia-Nicolas et al., 2023). As might be expected, viral load and pDC abundance showed positive correlations, with pDC abundance lower for the BA.5+MP group across all viral loads (Fig. 3b). pDC are dominant producers of type I interferons (IFNs) during SARS-CoV-2 infection, with low numbers of pDCs and low type I IFN levels generally associated with increased COVID-19 severity (Contoli et al., 2021; Van der Sluis et al., 2022).

Consistent with reduced abundance of pDC in the BA.5+MP group, was a series of type I IFN IPA USR annotations with high negative z scores (Fig. 3a, Type I IFNs; Supplementary Table 5). Reduction in type I IFN signatures was also evident from GSEAs using MSigDB gene sets (Supplementary Table 5). Despite significantly lower type I IFN signatures in the BA.5+MP group, this group did not have significantly higher viral titres or higher levels of viral RNA (Fig. 2d,e). This may reflect both the ability of SARS-CoV-2 to inhibit type I IFN responses, and compensation via other innate anti-viral pathways (Bastard et al., 2022; Rawle et al., 2021).

Either way, appropriate type I IFN responses are likely a key component of protective inflammation (Contoli et al., 2021; Mangalmurti et al., 2020). Consistent with reduced phagocytosis, M1 macrophages, pDCs and type I IFNs, was the general overall reduction in innate sensing (Fig. 3a, Innate sensing).

SpacialDecon, and GSEAs using MSigBD and Blood Transcription modules (BTMs), identified significantly reduced NK cell signatures in the BA.5+MP group (Fig. 3a, NK cells; Supplementary Table 5), with low early type I IFN levels linked to reduced NK activity during viral infections generally (Paolini et al., 2015) and likely also SARS-CoV-2 infections (Acharya et al., 2020). NK cells have the capacity to exert important early innate antiviral activities; however, SARS-CoV-2 shows a remarkable ability to evade this arm of the immune system (Lee et al., 2023). NK and NKT cells are important sources of early IFNγ (Paolini et al., 2015), thus the high negative z score for the IPA USR annotation for IFNγ (Fig. 3a, Type I IFN; Supplementary Table 5) is consistent with reduced abundance of NK and NKT cells.

The general reductions in innate immune signatures for BA.5+MP vs. BA.5-MP (Fig. 3, Innate sensing, Type I IFNs, NK cells) is likely to be responsible for the negative z scores for a series of IPA USR pro-inflammatory cytokine annotations (Fig. 3a, Proinflammatory cytokines; Supplementary Table 5). Overall these data illustrate that MPs can significantly ameliorate SARS-CoV-2-mediated activation of innate immune responses in the lung at 2 dpi.

### 3.7. Haematopoietic stem cell progenitors are elevated by MPs 2 dpi

Three cytokine signatures associated with haematopoiesis were identified as down-regulated in the IPA USR analysis; CSF1/M-CSF, CSF2/GM-CSF and IL3 (Fig. 3, Haematopoiesis; Supplementary Table 5). Cell deconvolution using SpatialDecon and the Nanostring gene lists also identified a higher abundance of haematopoietic stem and progenitor cells (HSPC) in lungs 2 dpi in the BA.5+MP group (Fig. 3a, Stem.Prog; Supplementary Table 5). M-CSF, GM-CSF and IL3 (as well as IFNα, IFNγ and IL1, and TLR) are some of the key cytokine signalling pathways that promote HSPC differentiation (Chavakis et al., 2019). These analyses thus suggest an accumulation of undifferentiated HSPC due to reduced differentiation into myeloid and/or lymphoid precursors,

Dysregulated emergency myelopoiesis and immature myeloid cells are associated with poor outcomes in COVID-19 patients (Romo-Rodriguez et al., 2023; Schulte-Schrepping et al., 2020; Townsend et al., 2022), although these are observations from peripheral blood during COVID-19 disease, rather than from lungs early post infection. Nevertheless, these data (Fig. 3a, Haematopoiesis & Haematopoietic stem and progenitor cells) suggest MPs in the lung suppress haematopoiesis, likely as a result of the overall reduction in the proinflammatory milieu. To clarify the terminology used in these annotations; haematopoiesis includes myelopoiesis, lymphopoiesis and erythropoiesis, whereas leukopoiesis encompasses myelopoiesis and lymphopoiesis.

### 3.8. Some anti-inflammatory signatures increased by MPs at 2 dpi

IPA analysis of BA.5+MP vs. BA.5-MP indicated some signatures that are usually associated with anti-inflammatory activity (Fig. 3a, Anti-inflammatory). The high USR z score for TRIM24 (Supplementary Table 5) argues that, although M1 macrophages are down, M2 macrophages are not increased, given that TRIM24 expression is suppressed in M2 macrophages (Yu et al., 2019).

Interstitial macrophages can have immunosuppressive properties (Schyns et al., 2019; Ural et al., 2020; Vanneste et al., 2023); however, although identified as down in Fig. 1 they were not identified as increased for BA.5+MP vs. BA.5-MP. Conceivably, these anti-inflammatory signatures arise from the higher abundance of alveolar macrophages that are not M1.

### 3.9 MPs promote some proinflammatory signatures at 6 dpi

RNA-Seq was also undertaken for BA.5+MP vs. BA.5-MP at 6 dpi (the day of peak pathology in this model) with reads again aligned to the mouse genome. Using the same filter (q<0.05) 528 DEGs were identified and were analysed as above (Supplementary Fig. 6). A number of proinflammatory cytokine IPA USR signatures were up-regulated (Fig. 4, Proinflammatory cytokines), all are associated with increased COVID-19 severity; TNF (Kokkotis et al., 2022; Song et al., 2023), IL1B (Labzin et al., 2023), IL1A (Sanchez-de Prada et al., 2022), OSM (Arunachalam et al., 2020), IL17A (Yin et al., 2023a), and IL6 (Avdeev et al., 2021). Upregulated activity of the transcription factor EPHAS is also associated with pulmonary inflammatory responses in lethal COVID-19 (Lopez-Cortes et al., 2022). The anti-inflammatory interleukin 1 receptor antagonist (IL1RN) signature was down-regulated, with reduced IL1RN recently associated with increased COVID-19 severity (Mukundan et al., 2023).

**Figure 4.**
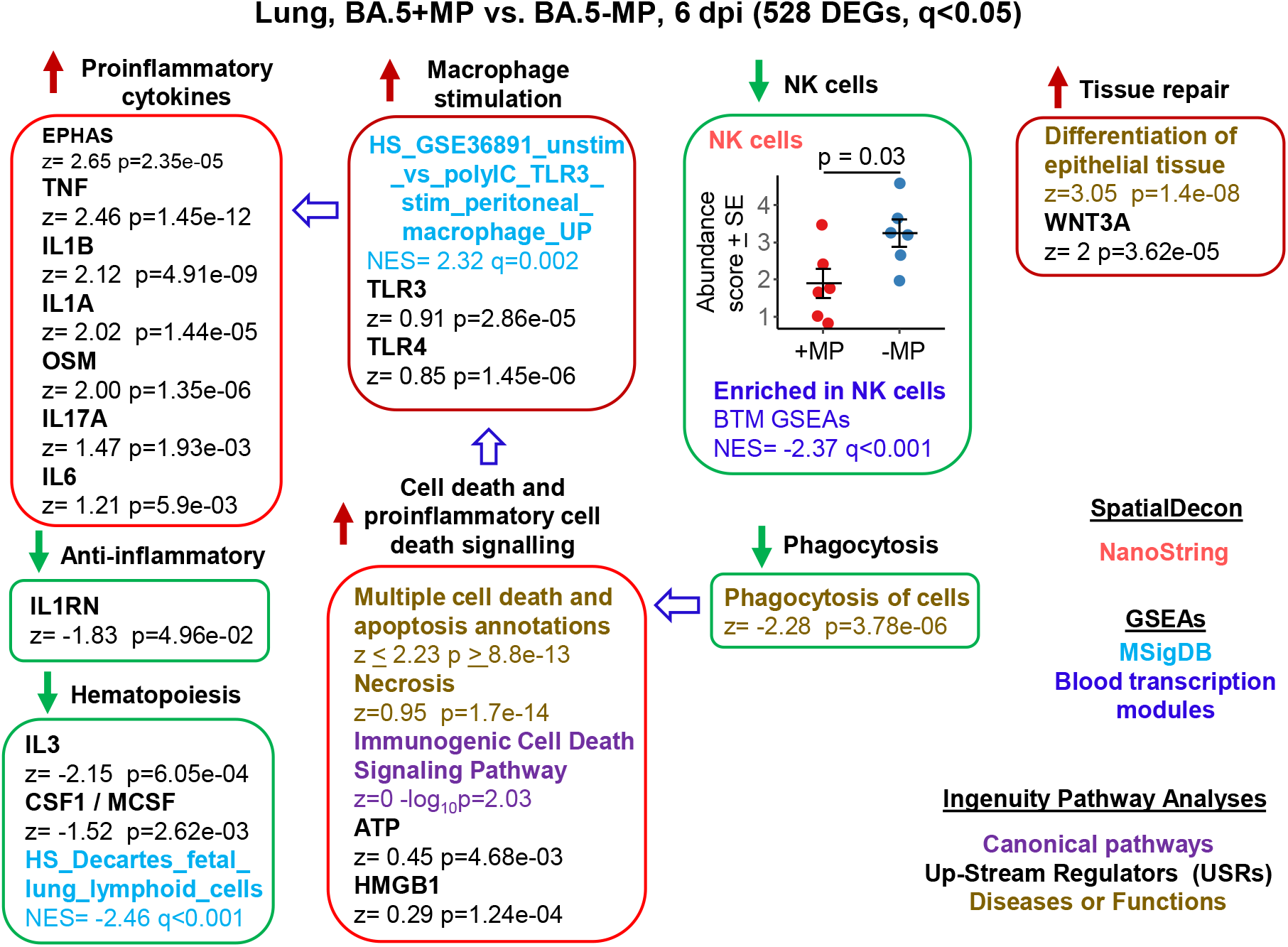
Bioinformatic analyses of RNA-Seq data for BA.5+MP vs. BA.5-MP at 6 dpi. **a** mACE-hACE2 mice were inoculated with BA.5+MP or with BA.5-MP and at 6 dpi lungs were harvested and analysed by RNA-Seq; 528 DEGs were identified. Bioinformatic analyses as in Fig. 3. Full data sets are provided in Supplementary Table 6.

The mechanisms responsible for these changes are not overtly apparent from the bioinformatic analyses; however, GSEAs using MSigDB gene sets provided a significant macrophage annotation with high NES for TLR3 stimulation, with an IPA USR annotation for TLR3 signalling also identified, albeit with a relatively low z score (Fig. 4, Macrophage stimulation). Viral double-stranded RNA is a probable ligand for TLR3 (Ullah et al., 2023). The reduced phagocytosis (Fig. 4, Phagocytosis), and the increased proinflammatory responses (Fig. 4, Proinflammatory cytokines), might suggest increased secondary necrosis due to a reduction in the phagocytosis of apoptotic cells (efferocytosis) (Ge et al., 2022; Razi et al., 2023; Sachet et al., 2017). Multiple IPA Disease & Functions annotations suggest an increase in apoptosis signatures (Supplementary Table 6), with secondary necrosis known to be proinflammatory via secretion of a number of mediators such as HMGB1 and ATP (Sachet et al., 2017). These latter mediators were identified as IPA USRs, although again with relatively low z scores (Fig. 4, Cell death and proinflammatory cell death signalling).

NK cell and haematopoiesis signatures remained, as at 2 dpi, down-regulated by MPs (Fig. 4). Tissue repair signatures were also identified (Fig. 4, Tissue repair) as might be expected by 6 dpi, when virus has been largely cleared.

### 3.10. MPs promote a ‘cytokine release syndrome’ profile in BA.5 infected lungs at 6 dpi

Human lung RNA-Seq data for severe lethal COVID-19 infections was recently provided, with two signatures described, a ‘Classical signature’ and a ‘cytokine release syndrome’ (CRS) signature (Budhraja et al., 2022). We re-derived two DEG lists from the fastq files deposited for this study (Supplementary Table 7; PRJNA1036279), with the lists then analysed by IPA. Significant cytokine USR z scores provided by this analysis were compared with the z scores of significant cytokine USRs identified for BA.5+MP vs. BA.5-MP (Fig. 5 a, b; Supplementary Table 7). A significant correlation emerged for CRS (Fig. 5a). In both human and mouse data sets, TNF, IL1A, IL1B, OSM, IL6 and IL17 signatures were prominently up-regulated, and CSF1, IL3 and EPO were prominently down-regulated (Fig. 5a, pink shading). These cytokines and their association with severe COVID-19 are described above for Fig. 4. Treatment with EPO (Maufak et al., 2022) and CSF1 (De Ponti et al., 2022; Mirchandani et al., 2022) have been considered for COVID-19, with low IL-3 levels associated with increased severity (Bénard et al., 2021).

**Figure 5.**
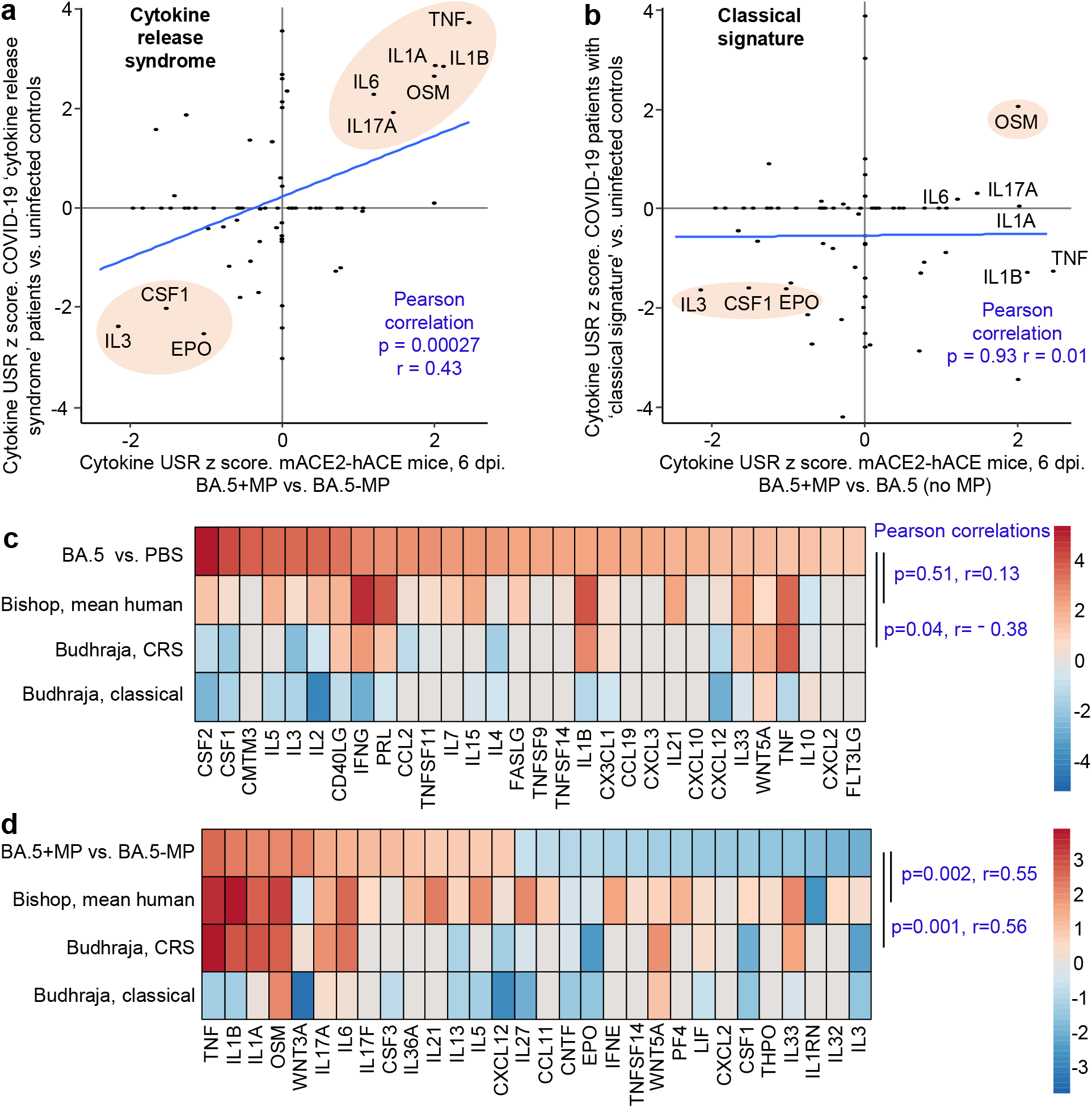
IPA cytokine USR z score correlations between human studies and BA.5+MP vs. BA.5-MP 6 dpi. **a, b** IPA Cytokine USR z scores from the IPA analysis of the 528 DEGs described in Fig. 4 were plotted against IPA Cytokine USR z scores from the IPA analysis of DEGs generated from fastq files obtained from NCBI SRA Bioproject PRJNA761132 (Budhraja et al., 2022) (Supplementary Table 7). Two distinct patterns were described for severe COVID patients, ‘Cytokine release syndrome’ and ‘Classical signature’; Pearson correlation significance (p) and correlation coefficient (r) are provided for both. Pink shading shows USRs dominant (high z scores) in both human and mouse studies. **c** The top 30 cytokine USRs by absolute z-scores for BA.5 vs. PBS are compared to the human lung cytokine USR z scores described in Bishop et al., 2022 (derived from 4 studies of SARS-CoV-2 infected vs. uninfected), and the cytokine USR z scores for ‘Cytokine release syndrome’ and ‘Classical signature’ described in Budhraja et al., 2022 (same as ‘a & b’ above). **d** The top 30 cytokine USRs by absolute z-scores for BA.5+MP vs. BA.5-MP are compared as for ‘c’. Pearson correlation significance (p) and correlation coefficient (r) are provided.

Budhraja et al., 2022 also described a “Classical signature” as the more common pattern for lethal COVID-19 (Budhraja et al., 2022); a similar IPA analysis indicated no correlation when cytokine USR z scores for the ‘Classical signature’ were compared with cytokine USRs z scores identified for BA.5+MP vs. BA.5-MP (Fig. 5b).

Another way of representing such data is by heat maps (Bishop et al., 2022). When the top 30 cytokine USRs (by absolute z-score) for BA.5 vs. PBS were ranked, and shown next to the aforementioned z-scores for CRS and the Classical signature, there were no significant positive correlations (Fig. 5c, Pearson correlations). Concordance was also poor for comparisons with the mean cytokine USR z scores previously generated from 4 human studies of SARS-CoV-2 infected lung tissues (Bishop et al., 2022) (Fig. 5c, Bishop, mean human). These data (Fig. 5c) likely reflect the relatively mild disease seen in the BA.5 mACE2-hACE2 mouse model.

When the top 30 cytokine USRs for BA.5+MP vs. BA.5-MP were ranked and compared with z scores from CRS in COVID-19 infected patients and the Bishop et al., 2022 study, significant positive correlations did emerge (Fig. 5d) . As in Fig. 5b, there was no significant correlation with the classical signature (Fig. 5d).

Taken together these analyses suggest that MPs in the SARS-CoV-2-infected lung pushed the cytokine signatures towards a CRS profile. However, MPs did not induce overt clinical disease in the BA.5-infected mice, nor were we able to detect significant histological changes in lung sections (Supplementary Fig. 2b, Supplementary Fig. 4).

### 3.11 MPs promote expression of stress response genes during SARS-CoV-2 infection

At the top of the DEG lists for BA.5+MP vs. BA.5-MP are a number of genes that are associated with stress, and genes that have previously been identified as being induced after smoke or diesel particle inhalation, or in other lung diseases/disease models (Table 1).

**Table 1.** Many top up-regulated DEGs are associated with stress responses and are also identified in other lung diseases or disease models. Top up-regulated DEGs on 2 and 6 dpi for BA.5+MP vs BA.5-MP (Supplementary Table 5; Supplementary Table 6). Log_2_FC are shown, with “-“ indicated if the gene is not a DEG on that day. The numbers in brackets (e.g. (1)) represent the position in the DEG list sorted by FC, i.e. Hspa1b is the most up-regulated DEG 6 dpi. ^1^ Genes that were synergistically induced for BA.5+MP vs BA.5-MP such that the cpm ratios (BA.5+MP/(BA.5 plus +MP) were >1 (Supplementary Table S8). ^2^No neutrophil-associated signatures were identified in any of the bioinformatic analyses, suggesting that in this setting this activity did not manifest.

| Gene | Log<sub>2</sub>FC<br>2 dpi | Log<sub>2</sub>FC<br>6 dpi | Function | Regulation/activity in lungs |
| --- | --- | --- | --- | --- |
| Hspa1b | - | 2.64<sup>1</sup><br>(1) | Stress-induced transcription chaperones. Cytoprotection (Hsp70 family members) | Biomarker for bronchopulmonary dysplasia (Evgen'ev et al., 2023). Upregulated in ARDS model (Rico-Llanos et al., 2022). Promotes epithelial cell repair (Duan et al., 2014) |
| Hspa1a | - | 2.46<sup>1</sup><br>(2) |  |  |
| Cxcl5 | - | 2.28<sup>1</sup><br>(3) | Chemokine. Tissue remodelling, neutrophil<sup>2</sup> recruitment. | Correlates with lung function decline in COPD patients and mouse smoking model (Chen et al., 2019) |
| Nr4a1, nuclear receptor subfamily 4 group A member 1 | 1.52<sup>1</sup><br>(1) | 1.77<br>(5) | Nuclear receptor with transcription activator activity. | Induced by acrolein (smoke chemical) in A529 cells (Thompson et al., 2008). |
| Ccn1 | 1.43<sup>1</sup><br>(2) | 1.33<br>(6) | Matricellular protein; inflammation, tissue repair. | Associated with ARDS severity (Morrell et al., 2020) |
| Cxcl1 | - | 1.09<br>(9) | Chemokine. Wound repair, neutrophil<sup>2</sup> recruitment. | Induced by diesel exhaust in mouse model (Kim et al., 2020) |
| Egr3 | 1.03<br>(10) | 1.07<br>(10) | Immediate early stress response gene | Upregulated by acrolein (Thompson et al., 2008). Upregulated in lungs from COPD patients (Chen et al., 2021) |
| Atf3<br>Activating transcription factor 3 | 0.54<br>(51) | 0.94<br>(14) | A master regulator of stress responses | Induced in lungs by particulate matter (Yan et al., 2018). Key role in lung regeneration (Niethamer et al., 2023). |
| Klf2, Krüppel-like Factor 2 | 0.82<br>(14) | 0.89<br>(16) | Transcription factor, endothelial integrity, shear stress induced, (Sangwung et al., 2017) | Induced by air pollution (twin study) (Madaniyazi et al., 2020). Ameliorates lung injury, animal models (Huang et al., 2017) |

Interestingly, the DEG overlap was low for up-regulated genes for BA.5+MP vs. BA.5-MP when compared with up-regulated genes for +MP vs. PBS (i.e. -MP) (Supplementary Fig. 5). Thus on their own MPs induced a largely different set of genes, when compared with MPs in a SARS-CoV-2 infection setting. This observation supports the contention that the detrimental activity of MPs might best be observed in the dysregulation of inflammation during a specific disease process (Rawle et al., 2022), rather than as an imposition of a fixed MP-specific response. To gain insights into the genes that might underpin this dysregulation, we identified genes that were synergistically induced (i.e. genes for which the cpm for +MP alone plus BA.5 alone > +MP+BA.5). Five genes emerged Hspa1a, Hspa1b, Cxcl5, Nr4a1 and Ccn1 (Table 1, Supplementary Table 8). The first 2 are members of the Hsp70 family, with Hsp70 induction well described in the MP literature (see below).

## 4. Discussion

We show herein in a mild disease model of SARS-CoV-2 infection and disease (omicron BA.5 infection of mACE2-hACE2 mice) that MP inoculation into the lungs dysregulated the inflammatory responses against the virus. Innate proinflammatory immune signatures were generally depressed at 2 dpi, but were elevated at 6 dpi. The profile of cytokine signatures in the lungs at 6 dpi showed significant correlation with the ‘cytokine release syndrome”, which is a potentially lethal manifestation of severe COVID-19 (Budhraja et al., 2022; Ghosh et al., 2023). However, despite this modulation, MP inoculation did not induce overt clinical or histologically-detectable changes in infected mice, suggesting the MP-mediated influences on SARS-CoV-2-mediated disease were clinically mild.

It is tempting to speculate that the ability of MPs to block efferocytosis via binding to the efferocytosis receptor Tim4 (Kuroiwa et al., 2023) provides a basis for understanding, at least some of the observations presented herein. Specifically, the clearly depressed phagocytosis signatures, the identification of multiple annotations associated with modulated macrophage responses (with macrophages the key mediators of efferocytosis), and the overall lack of detectable changes to adaptive immune responses, might be viewed as consistent with a role for dysregulated phagocytosis/efferocytosis (Ge et al., 2022; Razi et al., 2023). Reduced phagocytosis of SARS-CoV-2 infected cells at 2 dpi might reduce M1 macrophage activation, subsequent pDC activation (Garcia-Nicolas et al., 2023), and the ensuing cytokine responses (Severa et al., 2021; Van der Sluis et al., 2022). Less phagocytosis at the peak of infection might also lead to more secondary necrosis and/or necroptosis of SARS-CoV-2 infected cells, thereby promoting inflammation at 6 dpi (Fredman et al., 2023; Li et al., 2020).

Inhibiting phagocytosis/efferocytosis may not be the only mechanism in play, as MPs alone provided some stress, damage and pro-inflammatory signatures (Fig. 1), which may also influence the SARS-CoV-2-associated innate responses in the lung. The top DEGs for BA.5+MP vs. BA.5-MP support this contention as some of these DEGs have also been identified in studies of smoke or diesel particle inhalation (Table 1). The top DEGs for 6 dpi were heat shock protein 70 (Hsp70) family members, Hspa1a and Hspa1b, with these genes also synergistically induced (Table 1; Supplementary Table 8). Hsp70 up-regulation is reported in a range of MP exposure settings including mussels (Detree et al., 2018), goldfish (Abarghouei et al., 2021) and *Daphnia* (Yin et al., 2023b). Hsp70 stress responses are involved in a vast range of pathologies (Nunes et al., 2023) and play a role in *inter alia* cytoprotection against environmental challenges (Evgen’ev et al., 2023) including SARS-CoV-2 infection (Rébé et al., 2022). The top cytokine USR by z score for +MP vs. PBS was IL-1β, a cytokine matured and released via inflammasome activation (Fig. 1). Our observation support a recent speculation that MPs might activate the inflammasome (Alijagic et al., 2023), with IL-1β also a dominant signature at 6 dpi for BA.5+MP vs. BA.5-MP. Although many reports suggest MPs induce ROS (Hu et al., 2020; Subaramaniyam et al., 2023), we did not identify an increase in ROS as a major consequence of MP exposure, with the MPs used in this study being free of azide (Pikuda et al., 2019).

Nr4a1 (Table 1) was recently identified as a marker of a subset of group 2 innate lymphoid cells (ILCs) (Xu et al., 2023), with ILCs implicated in our previous study of MPs and a viral arthritis model (Rawle et al., 2022). However, we have been unable to find a compelling signature that implicates ILCs as important players in the current setting. Nr4a1-dependent CD16.2^+^ monocytes have been implicated as potential precursors of CD206^−^ interstitial macrophages (Schyns et al., 2019; Strickland et al., 2023; Vanneste et al., 2023). However, changes in interstitial macrophages were identified for +MP vs. PBS (Fig. 1), where Nr4a1 was not a DEG, but interstitial macrophages were not identified for BA.5+MP vs. BA.5-MP (Figs. 3 and 4) where Nr4a1 was a top DEG (Table 1). Conceivably, the up-regulation of this gene is associated with lung epithelial cells, where it has been identified as a novel allergy-associated gene (Zhu et al., 2022). MP-induced changes to the lung microbiota (Zha et al., 2023) might also provide a potential mechanism for the modulation of innate responses; however, our metagenomics analysis failed to identify any changes in the microbiome (Supplementary Fig. 6).

This study has a number of limitations; we have not investigated the activity of MPs with different shapes, sizes or compositions (Rozman et al., 2022); however, such considerations give rise to an unworkably large number of experimental variables. We have also not examined longer exposure times or different exposure doses, and we have also only examined the consequences of MPs in one mild mouse model of COVID-19. However, the mild MP-associated transcriptional changes (Supplementary Table 1) might be difficult to detect in a more severe model of COVID-19 where expression of a large number of genes are substantially changed.

In summary, we provide herein evidence that MPs in the lung can dysregulate innate inflammatory transcriptional responses to SARS-CoV-2 in a mouse model of mild COVID-19 infection. However, the dysregulation did not result in overt changes in disease or histopathology, suggesting MP-mediated changes were generally mild. To what extent such changes might influence COVID-19 disease severity at a population level may warrant investigation by perhaps comparing matched populations exposed to high (Liu et al., 2019b) and low levels (GGGI, 2022) of airborne MP pollution, although separating the influence of MP from other factors would represent a formidable challenge.

## 5. Conclusion

Considerable speculation regarding the health impacts of MP inhalation remains (Fiorito et al., 2022; Liaquat et al., 2022; Perera et al., 2023; Yang et al., 2022), with limited compelling *in vivo* data from humans and animal models. However, an emerging theme from such studies, including the current report, is that MPs can dysregulate and/or promote inflammatory processes in specific disease settings (Chen et al., 2023; Liu et al., 2022; Luo et al., 2022; Rawle et al., 2022; Taş et al., 2023; Yan et al., 2022c). However, pertinent insights can be developed only if animal models use realistic MP doses, although we need (i) better insights into what such doses actually are in different human populations, and (ii) standardised units for MP exposure (Liu et al., 2024) (e.g. µg/kg/d) so that studies can be compared. Avoidance of artefacts associated with preservatives such as azide is clearly also critical for future meaningful medical research in the MP space (Pikuda et al., 2019).

## CRediT authorship contribution statement

Conceptualization, A.S.; Methodology, C.R.B., D.J.R. and A.S.; Formal analysis, C.R.B., T. L., W.N.; Investigation, K.Y., T. L. and B.T.; Data curation, C.R.B., A.S.; Writing – original draft, A.S. Writing – review and editing, C.R.B.; Visualization, C.R.B., A.S.; Supervision, D.J.R., W.N. and A.S.; Project administration, A.S.; Funding acquisition, A.S. D.J.R.

## Data availability

All data is provided in the manuscript and accompanying supplementary files. Raw sequencing data (fastq files) generated for this publication for RNA-Seq have been deposited in the NCBI SRA, BioProject: PRJNA1036279 and are publicly available at the date of publication.

## Acknowledgements

The authors thank the following QIMRB staff; Dr. I. Anraku for management of the PC3 facility at QIMR Berghofer MRI, Dr. Viviana Lutzky for proof reading, Drs Clay Winterford and Crystal Chang for histology services, the animal house staff for mouse breeding and agistment, and Dr. Gunter Hartel for assistance with statistics.

