## Supplementary Figures for "Microplastics dysregulate innate immunity in the SARS-CoV-2 infected lung"

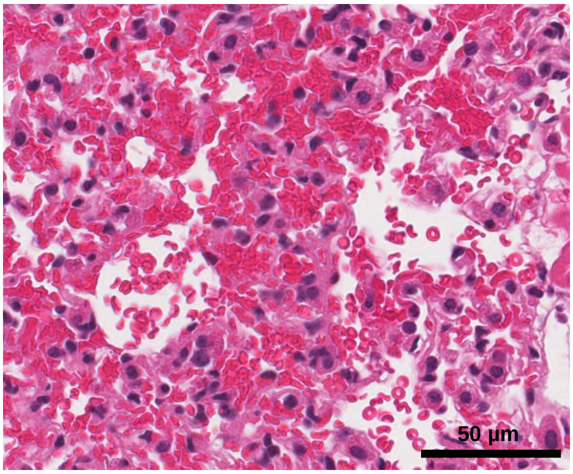


Focal alveolar haemorrhage

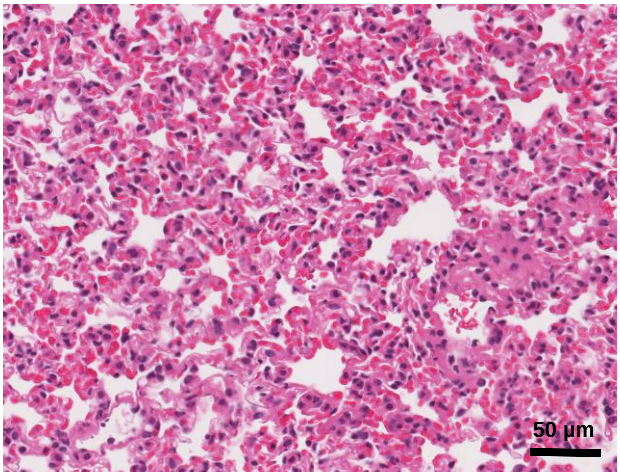


Lung consolidation, thickening of alveolar septa
(loss of alveolar airspaces)

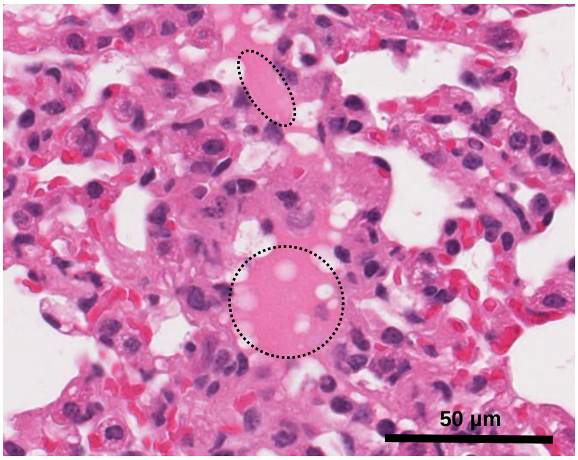


Alveolar edema

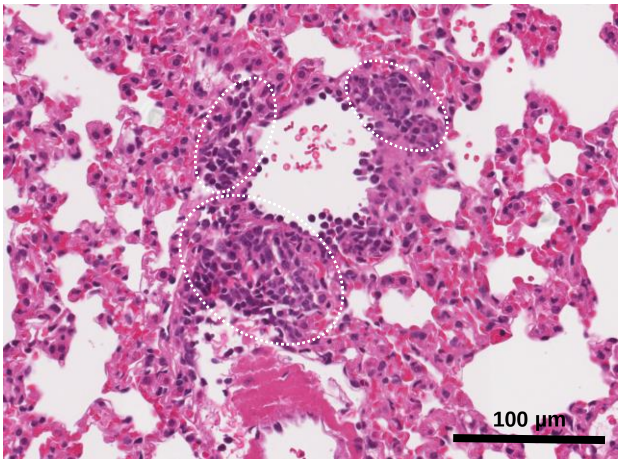


Focal leukocyte infiltration

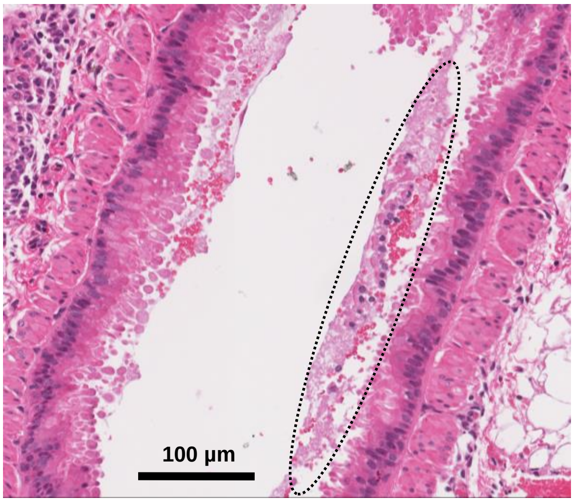


Serum and red blood cells in airways
(mild sloughing of bronchial epithelium)

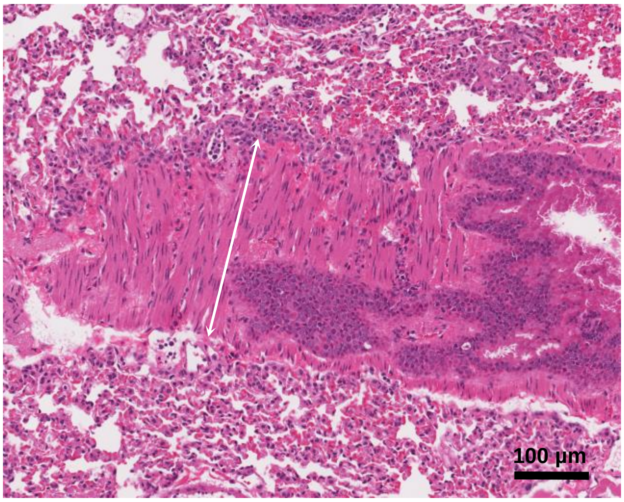


Smooth muscle hyperplasia/hypertrophy

**Supplementary Figure 1**. Images of H&E stained sections from lungs of mACE2-hACE mice infected with omicron BA.5, 6 dpi. Histopathological features described herein have been reported previously for mouse models of COVID-19.

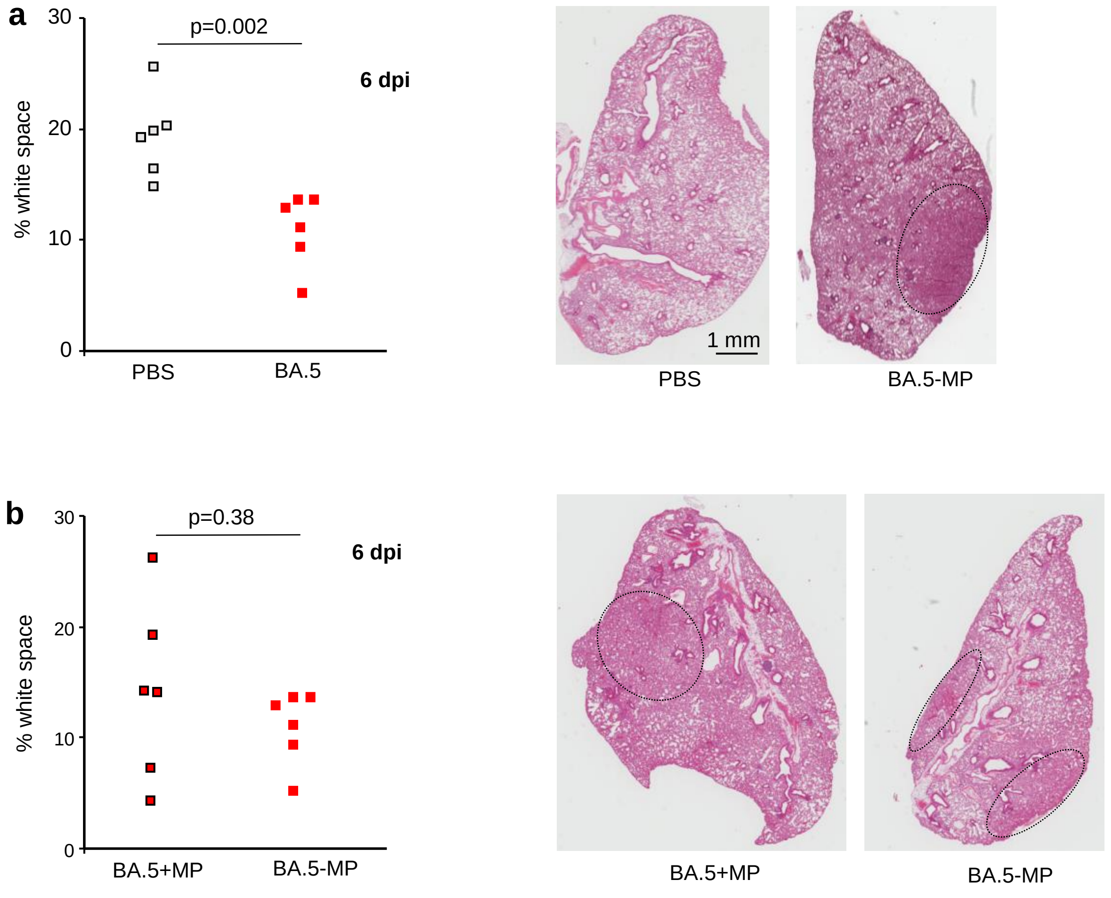


**Supplementary Figure 2**. **a** H&E stained sections of lungs of mACE2-hACE mice inoculated with PBS or infected with BA.5 (6 dpi), where analysed for white space; the area within the lung that remains unstained (white) as a percentage of the total lung area is shown for each mouse. Statistics by t test. H&E images show examples of whole lung sections stained with H&E; an area with overt loss of white space (lung consolidation) is indicated by the dotted oval. **b** As for “a” but comparing lungs receiving BA.5+MP vs. BA.5-MP (BA.5 with no MP).

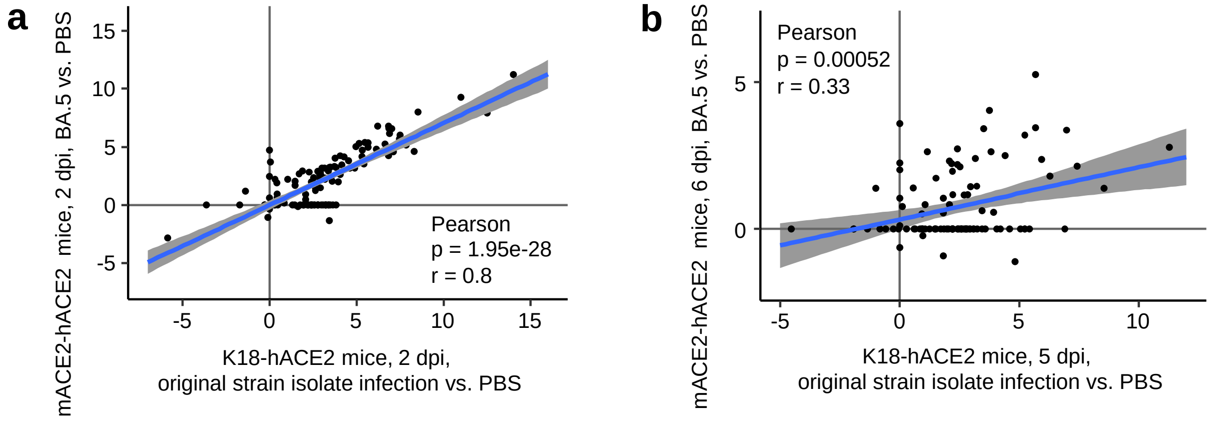


**Supplementary Figure 3**. **a** Pearson correlation of IPA USR z-score for Cytokine annotations for lungs at 2 dpi for the two indicated mouse models. **b** As for ‘a’ but at 5/6 dpi.

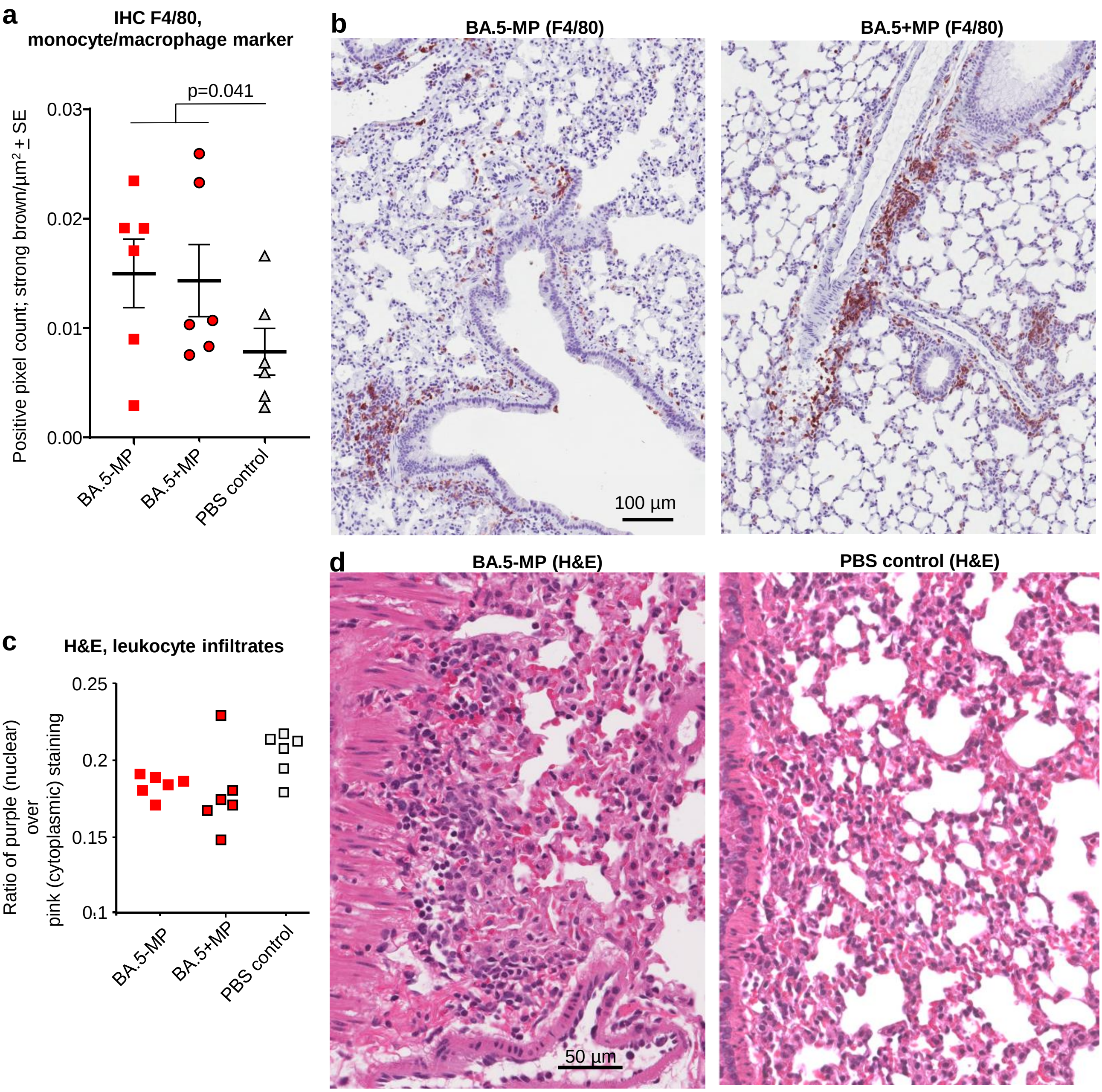


**Supplementary Figure 4**. **a** Positive pixel count analysis of IHC stained (F4/80) sections of lungs of mACE2-hACE mice infected with BA.5 (BA.5-MP), infected with BA.5 and inoculated with MP (BA.5+MP), or inoculated with PBS (PBS control). Lung were harvested on 6 dpi. The small increase in the infected groups reached significance if both infected groups were taken together (n=12), statistics by t test. **b** Examples of IHC staining in sections with high pixel counts. Positive cells (brown) are clearly visible, often in infiltrates adjacent to airways. (Counter staining with haematoxylin, light purple). **c** Positive pixel count analysis of H&E stained sections from the same mice as in "a". Purple (nuclear) over pink (cytoplasmic) staining is a broad (crude) measure of leukocyte infiltration, as leukocytes have a higher nuclear to cytoplasm ratio. However, no significant differences in cellular infiltrates were evident for BA.5+MP vs. BA.5-MP. **d** Examples of H&E staining. Infiltrates in infected mice were often diffuse and associated with lung consolidation & smooth muscle hyperplasia (left), such that the purple/pink ratios were not significantly higher even than controls (right). Other mouse models of COVID-19 have more severe lung disease and are associated with more abundant and less diffuse leukocyte infiltrates, such that purple/pink ratios are significantly higher in infected lungs when compared with controls (Rawle et al., 2022; Dumenil et al., 2022).

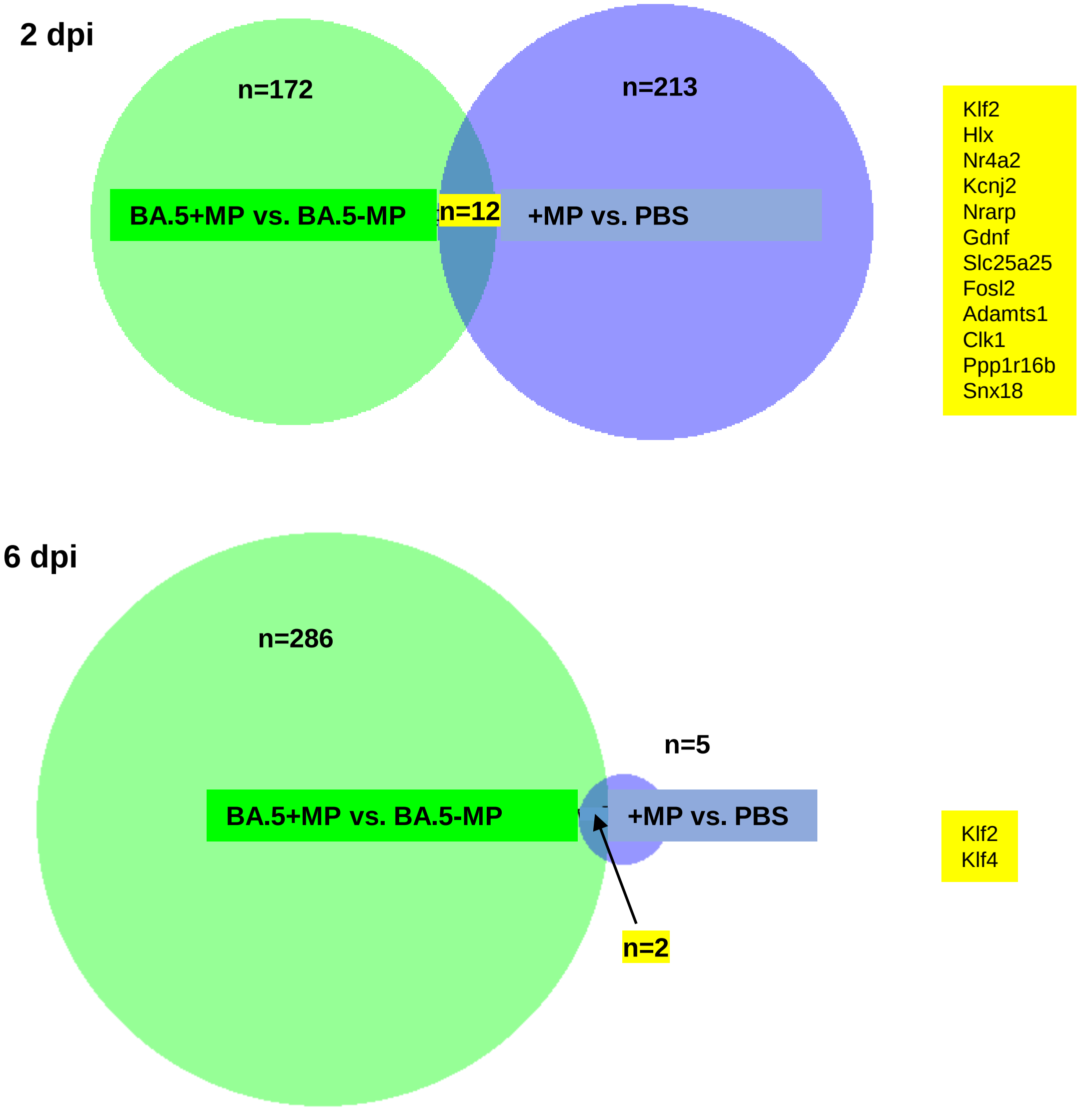


**Supplementary Figure 4.** Up-regulated DEGs from the indicated comparisons. Overlapping DEGs are indicated in yellow.

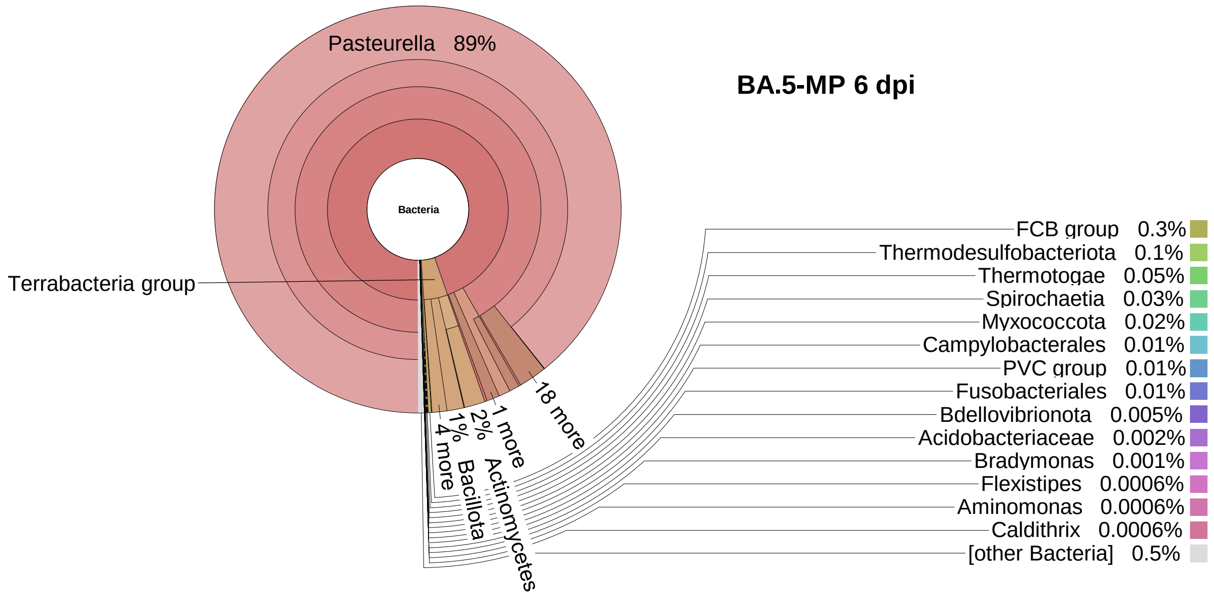


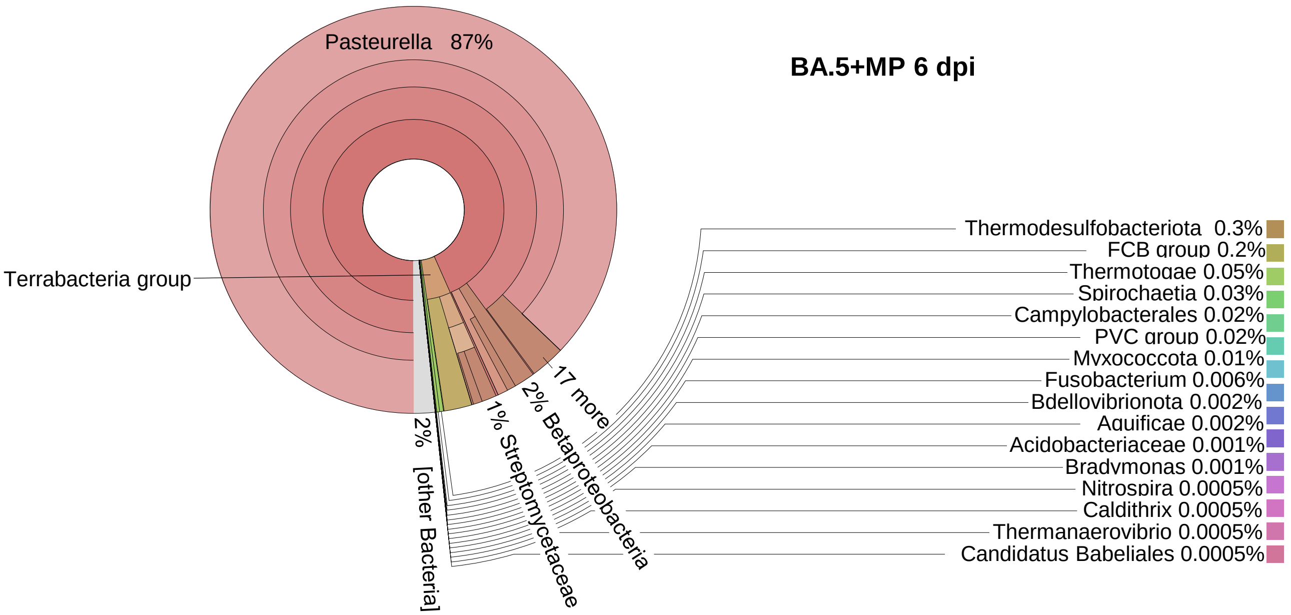


**Supplementary Figure 6.** Kraken metagenomic sequence classification undertaken as described (Hazlewood et al., 2021). Briefly, the unmapped reads were analysed using metagenomic sequence classification. Replicates were concatenated to produce one read file per treatment. Data was visualized using Krona, showing Bacteria (root).
